# Removal of Spo11 from meiotic DNA breaks *in vitro* but not *in vivo* by Tyrosyl DNA Phosphodiesterase 2

**DOI:** 10.1101/527333

**Authors:** Dominic Johnson, Rachal M Allison, Elda Cannavo, Petr Cejka, Matthew J Neale

**Affiliations:** Genome Damage and Stability Centre, School of Life Sciences, University of Sussex, Brighton UK; Institute for Research in Biomedicine, Università della Svizzera italiana, Bellinzona 6500, Switzerland; Department of Biology, Institute of Biochemistry, Eidgenössische Technische Hochschule (ETH) Zurich, Zurich 8093, Switzerland

## Abstract

Meiotic recombination events are initiated by DNA double-strand breaks (DSBs) created by the topoisomerase-like protein, Spo11. Similar to type-II topoisomerases, Spo11 becomes covalently linked to the 5′ ends generated on each side of the DSB. Whilst Spo11-oligos—the product of nucleolytic removal by Mre11—have been detected in a number of biological systems, the lifetime of the covalent Spo11-DSB precursor has not been systematically determined and may be subject to alternative processing reactions. Here we explore the activity of human Tyrosyl DNA Phosphodiesterase, TDP2, on Spo11-DSBs isolated from *S. cerevisiae* cells. We demonstrate that TDP2 can remove Spo11 from natural ssDNA-oligos, and dsDNA ends even when in the presence of excess competitor genomic DNA. Interestingly, TDP2-processed Spo11-DSBs are refractory to resection by Exo1, suggesting that ssDNA generated by Mre11 may be essential *in vivo* to facilitate resection-dependent HR at Spo11-DSBs even if TDP2 were active. Moreover, although TDP2 can remove Spo11 peptides *in vitro*, TDP2 was unable to remove Spo11 *in vivo*—unlike during the repair of topoisomerase-induced DNA lesions. These results suggest that Spo11-DNA, but not topoisomerase-DNA cleavage complexes, are inaccessible to the TDP2 enzyme, perhaps due to occlusion by higher order protein complexes resident at sites of meiotic recombination.

## INTRODUCTION

In most organisms, meiotic recombination is essential to create genetically diverse, haploid genomes suitable for sexual reproduction. In meiosis, homologous recombination (HR) events are initiated by DNA double-strand breaks (DSBs) created by Spo11, a topoisomerase-like enzyme most closely related to the catalytic ‘A’ subunit of the archael type-IIB Topoisomerase VI (1–3). TopoVI functions as an A_2_B_2_ heterotetramer, and the recent identification of the eukaryotic ‘B’ subunit, Top6BL, in mouse, plants and budding yeast indicates that the active form of Spo11 is similarly heterotetrameric (4, 5). Like other type-II and type-IIB topoisomerases, the Spo11 monomers become covalently attached via a phosphotyrosine bond to the 5′ ends generated on each side of the DSB during cleavage (6– 8). In wildtype *S. cerevisiae* cells, such covalent Spo11-DSBs are transient, being rapidly processed by the nuclease activity of the MRX/N complex (Mre11, Rad50 and Xrs2/Nbs1), which is stimulated by Sae2, the yeast orthologue of human CtIP (9–14). This coordinated reaction generates a covalent single-stranded Spo11-oligonucleotide complex, flanked by a nick or short ssDNA gap on the 5′-ending strand that enables the DSB to be subsequently channelled into homologous recombination repair via the onset of long-range ssDNA resection catalysed by Exo1 (9, 15–17). Spo11-oligo complexes are formed in an evolutionarily divergent yeast, *Schizosaccharomyces pombe* (18), and are also detectable in mouse spermatocytes (10, 19), suggesting that nucleolytic release is the default mode of Spo11 removal in both unicellular and multicellular eukaryotes.

However, whilst the product of this nucleolytic reaction—Spo11-oligos—have been detected in numerous biological systems, the lifetime of the covalent DSB precursor has not been systematically determined and may be subject to alternative processing reactions. For example, in *S. pombe*—and unlike in *S. cerevisiae* (2)—covalent Spo11-DSBs are readily detected in wildtype cells (20), indicating that DSB formation and the initiation of repair can be temporally separated. Moreover, inactivation of the canonical Spo11 release reaction in *C. elegans* (via mutation of COM-1, the CtIP/Sae2 orthologue) results in repair of DSBs by the non-homologous endjoining (NHEJ) pathway and aberrant chromosome segregation (21, 22). These observations suggest that delayed onset of meiotic DSB resection may be especially problematic for meiotic cells, and further suggest there must be a COM-1-independent mechanism to remove Spo11 in *C. elegans*, such that the DSBs can be ligated using the end-joining apparatus.

In vegetative cells, the usually benign and transient protein-linked DNA breaks created by Topoisomerase I (Top1) and II (Top2) as part of their reversible catalytic cycle may become a stable and toxic covalent-DNA intermediates (23). This conversion is exploited in chemotherapy by the use of drugs such as camptothecin and etoposide, which stabilise the covalent cleavage-complex intermediate (Top1cc and Top2cc respectively), thereby sensitising cells that have a relatively greater reliance on topoisomerase activity compared to other tissue. Presumably because such topoisomerase failures are relatively common even in unchallenged cells, two classes of enzymes exist that are capable of handling such lesions: Tyrosine phosphodiesterase 1 (Tdp1), which primarily resolves the 3′ covalent phosphotyrosine links created by Top1 (24), and tyrosine phosphodiesterase 2 (TDP2, also named TTRAP), which primarily resolves 5′ phosphotyrosine links created by Top2 (25). Some enzymatic overlap between the activities and substrates of Tdp1 and TDP2 exists, suggesting some functional redundancy *in vivo* (25, 26). Nevertheless, the primary activity of Tdp1 activity aids the conversion of Top1 lesions into ligatable single-strand breaks (SSBs), whereas TDP2 converts Top2 lesions into DSB ends suitable for direct repair by the NHEJ pathway. The reaction product created by TDP2 contrasts with the 3′ single-stranded ends generated as a result of MRX/N activity, and which are suitable for HR (9, 15, 27). Interestingly, TDP2 is expressed in mouse and human testes tissue (28), leading to the possibility that TDP2 may be able to process a subset of Spo11-DSB intermediates.

Here we use ectopic expression and *in vitro* biochemistry to explore the activity of human TDP2 on Spo11-DSBs generated in *S. cerevisiae* meiotic cells. Collectively, our results indicate that while TDP2 is catalytically capable of removing Spo11 from both Spo11-oligos and Spo11-DSB ends *in vitro*, TDP2 is unable to do so *in vivo*. This is despite TDP2 functioning efficiently to suppress the sensitivity of *mre11* or *sae2* mutants to topoisomerase poisons. These results suggest that Spo11-DSBs, but not Topoisomerase-DSBs, are inaccessible to the TDP2 active site, perhaps due to occlusion by higher order protein complexes resident at sites of meiotic recombination. Moreover, we find that the 2 nt 5′ extensions revealed when Spo11 is removed from DSB ends by TDP2 *in vitro* are refractory to nucleolytic resection by Exo1, suggesting that even were TDP2 to act during meiotic DSB repair, it would nonetheless lead to DSBs being channelled away from the essential process of homologous recombination. Overall, our findings provide insight into the mechanisms involved in the resection and repair of protein-linked DSB ends created by topoisomerase-like enzymes such as Spo11, and how they may differ to the repair of ‘clean’-ended DNA breaks created by site-specific nucleases.

## RESULTS

### TDP2 removes Spo11 protein and peptides from single-stranded Spo11-oligo complexes

Meiotic recombination is initiated by DSBs created by the evolutionarily conserved Spo11 protein. As part of the catalytic reaction, Spo11 becomes covalently attached to the 5′ DSB ends via a phosphotyrosyl bond. In *M. musculus, S. cerevisiae*, and *S. pombe*, Spo11 is released from the DSB ends by the action of the evolutionarily conserved MRX/N nuclease complex in conjunction with Sae2/CtIP/Ctp1, generating covalent Spo11-oligo complexes (9, 10, 18, 19, 29). However, whether alternative mechanisms of Spo11 end-processing are possible has not been investigated.

To investigate whether Spo11-DSBs may also be a substrate for processing by TDP2—as are Top2-induced DSBs (25)—we took advantage of the *S. cerevisiae* meiotic system in which synchronised populations of cells undergo Spo11-DSB formation at hundreds of sites spread across the genome (termed Spo11 hotspots), but in which TDP2 is not naturally expressed.

We first tested for the ability of recombinant TDP2 to remove Spo11 covalently attached to ssDNA by enriching for the natural covalent Spo11-oligo complexes created by MRX and Sae2-dependent processing of meiotic Spo11-DSBs in wildtype cells (Fig 1a). Cells in mid meiotic prophase (4 hours after meiotic entry) were harvested, lysed, and Spo11-oligo complexes isolated by immunoprecipitation using antibodies raised against a C-terminal triple FLAG epitope (30).

**Figure 1:**
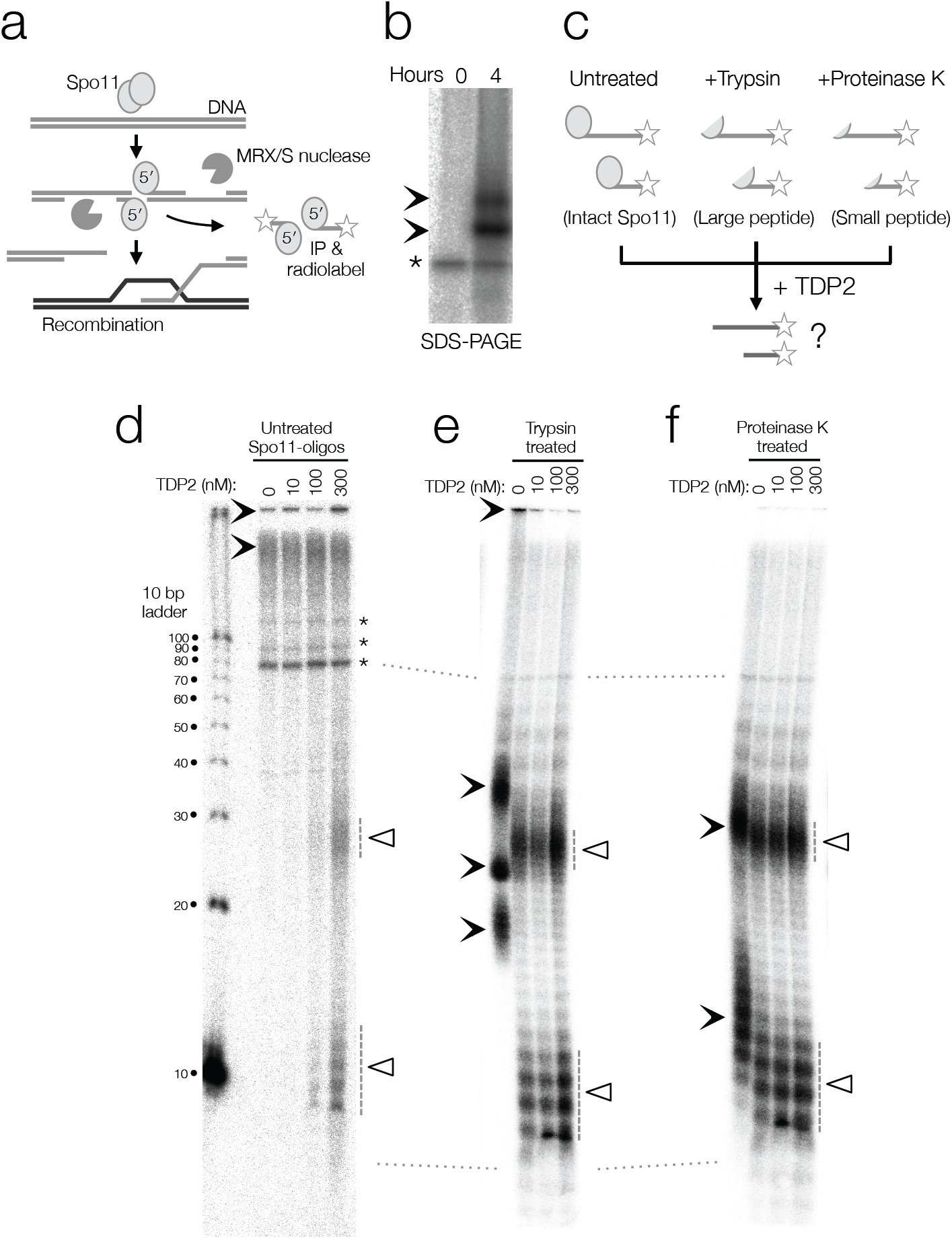
TDP2 can cleave the covalent phosphotyrosyl bond between Spo11 and a single-stranded oligonucleotide. a-b, Cartoon of the early stages of meiotic recombination. Spo11 generates DSBs and is released covalently attached to short oligonucleotides that are isolated by immunoprecipitation, and end-labelled at the 3′OH using alpha-^32^P dCTP and TdT, before resolving by SDS-PAGE (b). c-f, Schematic of the experiment (c). Labelled Spo11-oligo complexes without (d) or after treatment with Trypsin (e) or Proteinase K (f), were reacted with recombinant TDP2 at the indicated concentrations, then resolved on 18% polyacrylamide gel under denaturing conditions. Solid arrowheads indicate the migration position of the various Spo11-oligo species (prior to TDP2 treatment). Open triangles and dashed lines indicate the migration position of Spo11-oligos after Spo11-peptide removal by TDP2. Dotted connectors between gels indicate relative migration positions. Asterisks are a non-specific band.

Immunoprecipitates were washed and then the ssDNA component labelled at the terminal 3′ hydroxyl group using terminal deoxynucleotide transferase (TdT) and a radiolabelled nucleotide (dCTP). Under the conditions employed, this reaction results in 1-4 radiolabelled nucleotides being added to the 3′ OH end of the Spo11-oligo complex (D. Johnson and M. Neale unpub. obs), which can be resolved via SDS-PAGE (Fig 1b). Recombinant human TDP2 (25) was reacted with these “full-length” labelled Spo11-oligo complexes, or with complexes pre-treated with either trypsin or proteinase K (Fig 1c-f). Based on substrate specificity, trypsin is expected to leave twelve amino acid residues covalently attached to the oligo complex via the phosphotyrosine bond, whereas proteinase K treatment will degrade all but three amino acids in addition to the phosphotyrosine.

Following incubation with increasing concentrations of TDP2, reactions were halted by adding formamide gel-loading buffer, then resolved on a 15% polyacrylamide gel under DNA-denaturing conditions (Fig 1d-f). Under these conditions, untreated Spo11-oligo complexes fail to enter the gel and are often lost from the wells during gel handling and fixation (Fig 1d). By contrast, trypsin-treated molecules migrate as a heterogeneous smear, presumably due to significant electrophoretic retardation caused by the remaining amino acids bound to the DNA (Fig 1e). Proteinase K treated samples migrate more uniformly, as a pair of broad peaks, consistent with the previously characterised dual size classes of Spo11-oligo complex detected in *S. cerevisiae* (8-15 nt and 25-40 nt; Fig 1f; (9, 10)).

Consistent with experiments performed using synthetic phosphotyrosine ssDNA substrates (25) or partially proteolysed Top2-DNA covalent complexes (31), we observed robust activity of TDP2 against both the partially and fully proteolysed Spo11-oligo substrates (Fig 1e,f). In both cases, we observed the conversion of the radiolabelled signal to faster migrating forms consistent with cleavage of the 5′ phosphotyrosine bond and complete removal of any residual amino acids from the radiolabelled oligo.

Importantly, incubation of TDP2 with a radiolabelled oligononucleotide (i.e. without a covalent protein attachment) resulted in no change in electrophoretic migration (Fig S1a), confirming that the increased rate of migration observed above (Fig 1e,f) was not due to trace nuclease activity within the TDP2 preparation, but instead was due to the expected cleavage of the protein-DNA phosphotyrosyl bond.

Structural modelling of TDP2 indicates that the catalytic site is buried within the core of the protein, which any phosphotyrosine substrate would be required to thread into to be cleaved (32–34). Intriguingly, however, we also observed conversion of non-migrating to rapidly migrating oligo complexes when TDP2 was reacted with full-length Spo11-oligos, albeit only restricted to the reactions containing the higher enzyme concentrations (Fig 1d). These observations support the view that TDP2 can remove both full-length and Spo11 protein fragments from ssDNA ends, and that prior proteolytic degradation of the 5′-covalently linked protein stimulates the enzymatic activity of TDP2, similar to what has been observed for its activity on Top2-DNA intermediates (31).

### TDP2 removes Spo11 peptides from DSB ends

We next tested the ability of TDP2 to remove Spo11 from the naturally occurring double-stranded DNA (dsDNA) ends generated during meiosis. As introduced above, in wildtype *S. cerevisiae* cells, Spo11-DSBs are normally rapidly processed by MRX/Sae2 (liberating Spo11-oligo complexes) and then subjected to Exo1-dependent resection to create long 3′ ending ssDNA tails. Thus, to enrich for a population of covalent Spo11-DSB molecules to use as a substrate, and therefore to prevent such nucleolytic processing steps, we prepared genomic DNA samples from *sae2*Δ cells in mid meiotic prophase (4 hours after meiotic induction when DSBs are abundant). The resulting nucleic acid material is free from non-covalently attached protein contaminants but, due to the *SAE2* gene deletion, retains covalent Spo11-DSB intermediates (8, 35, 36). Samples were then incubated with and without TDP2 for up to 15 minutes, then subsequently reacted with lambda exonuclease—a 5′-3′ exonuclease with activity that depends on the presence of an unmodified 5′ phosphate group on the substrate (37). Post-treatment with lambda exonuclease thus provides a sensitive test for TDP2-dependent removal of the Spo11 moiety from the DSB end (Fig 2a).

**Figure 2:**
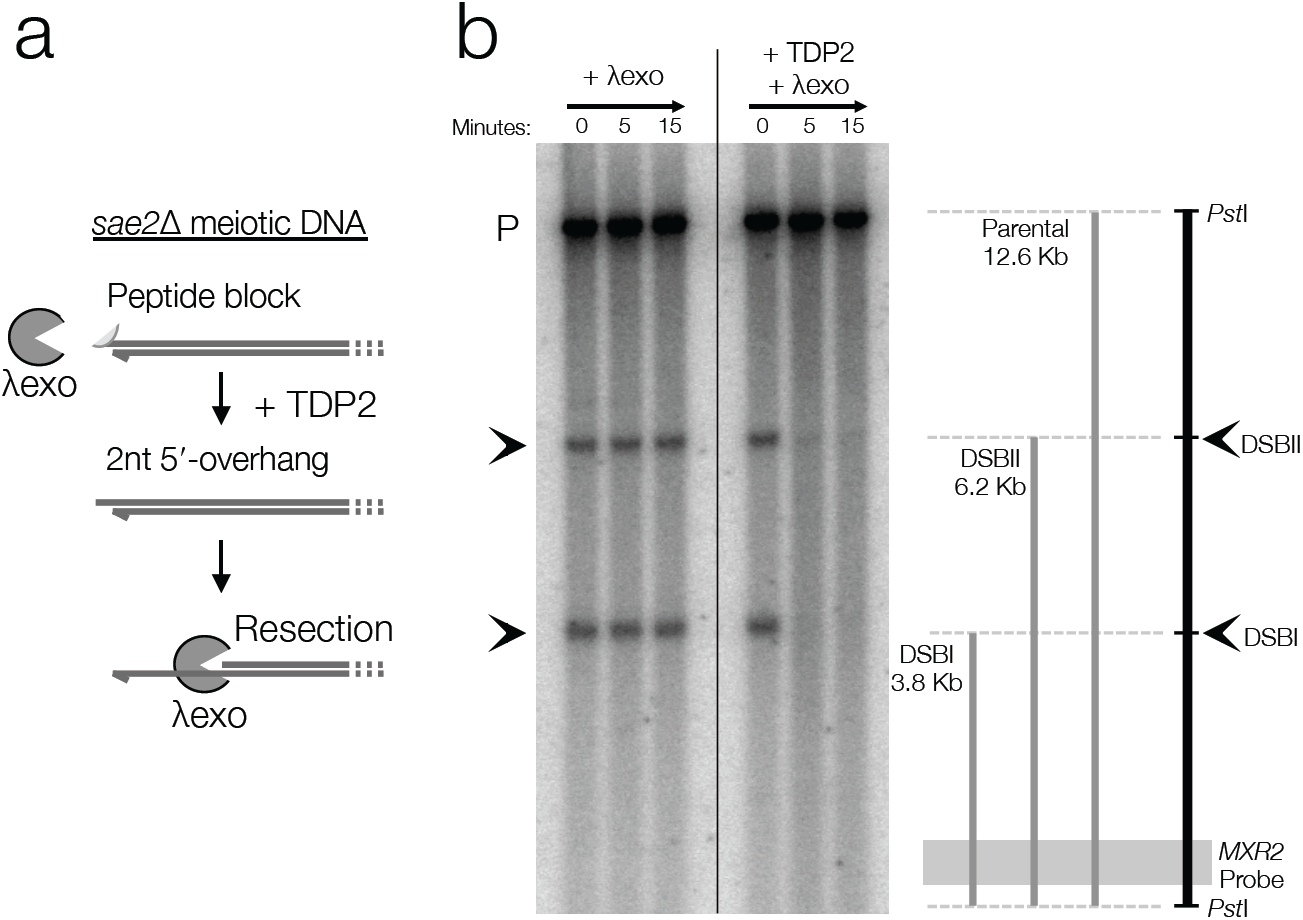
TDP2 can remove Spo11 peptides from the dsDNA end of natural Spo11-DSBs enabling 5′-3′ resection by lambda exonuclease. a, Schematic of experiment. b, Genomic DNA isolated from meiotic *sae2*Δ cells was treated with proteinase K and incubated with lambda exonuclease (λexo) without or with prior treatment with TDP2. DNA was purified and digested with *Pst*I, separated by electrophoresis on a 0.7% agarose gel, blotted to nylon membrane and hybridised with a probe (*MXR2* locus) close to the well-characterised meiotic recombination hotspot *HIS4*∷*LEU2*. The location of the two major Spo11-DSBs at this locus are marked with solid arrowheads. Disappearance of these signals following TDP2 and lambda exonuclease treatment is due to rapid exonucleolytic degradation from the uncapped Spo11-DSB ends.

To monitor the reactions, DNA was digested with *Pst*I, separated on an agarose gel, transferred to nylon membrane and hybridised with a radiolabelled DNA probe that recognises the *HIS4*∷*LEU2* meiotic recombination hotspot (38). Preparation in this way reveals three distinct bands, the upper most is the parental DNA molecule, and the lower smaller bands are indicative of DSBs arising at the two strong hotspots present at this genomic locus (Fig 2b). Treatment with lambda exonuclease alone resulted in no change in the migration of the DSB bands—as expected due to the attached Spo11 protein preventing 5′-3′ resection (Fig 2b). By contrast, pretreatment with TDP2 led to complete disappearance of the DSB bands—as expected if TDP2 was able to efficiently remove Spo11 from the DSB end creating a clean 5′ phosphorylated substrate on which lambda exonuclease can act (Fig 2b). Notably, the parental DNA band was stable under these conditions indicating that the exonuclease degradation was specific to the deprotected Spo11-DSB ends (Fig 2b). If we assume there to be approximately 160 DSBs (39) within each replicated *S. cerevisiae* diploid genome (∼50 Mbp) there will be approximately one covalent Spo11-DSB end for every ∼150 kb of genomic DNA. We conclude that recombinant TDP2 is able to efficiently remove Spo11 from dsDNA ends even in the presence of a large excess of dsDNA.

### Exo1 is unable to resect Spo11-DSBs that have been processed by TDP2

As described above, the endogenous exonuclease involved in 5′-3′ resection of Spo11-DSBs is Exo1 (9, 15, 17). We were thus interested to examine whether treatment of Spo11-DSBs with TDP2 could enable resection by recombinant Exo1, perhaps indicating a potential mechanism to bypass the requirement of MRX/N nuclease activity *in vivo*. To perform these experiments, we isolated genomic DNA from *sae2*Δ cells as described above, incubated with and without TDP2 for up to 15 minutes, then incubated with recombinant Exo1 rather than lambda exonuclease. Unlike treatment with lambda exonuclease, Exo1 was unable to initiate resection on Spo11-DSBs whether or not they had been processed by TDP2 (Fig 3a,b).

**Figure 3:**
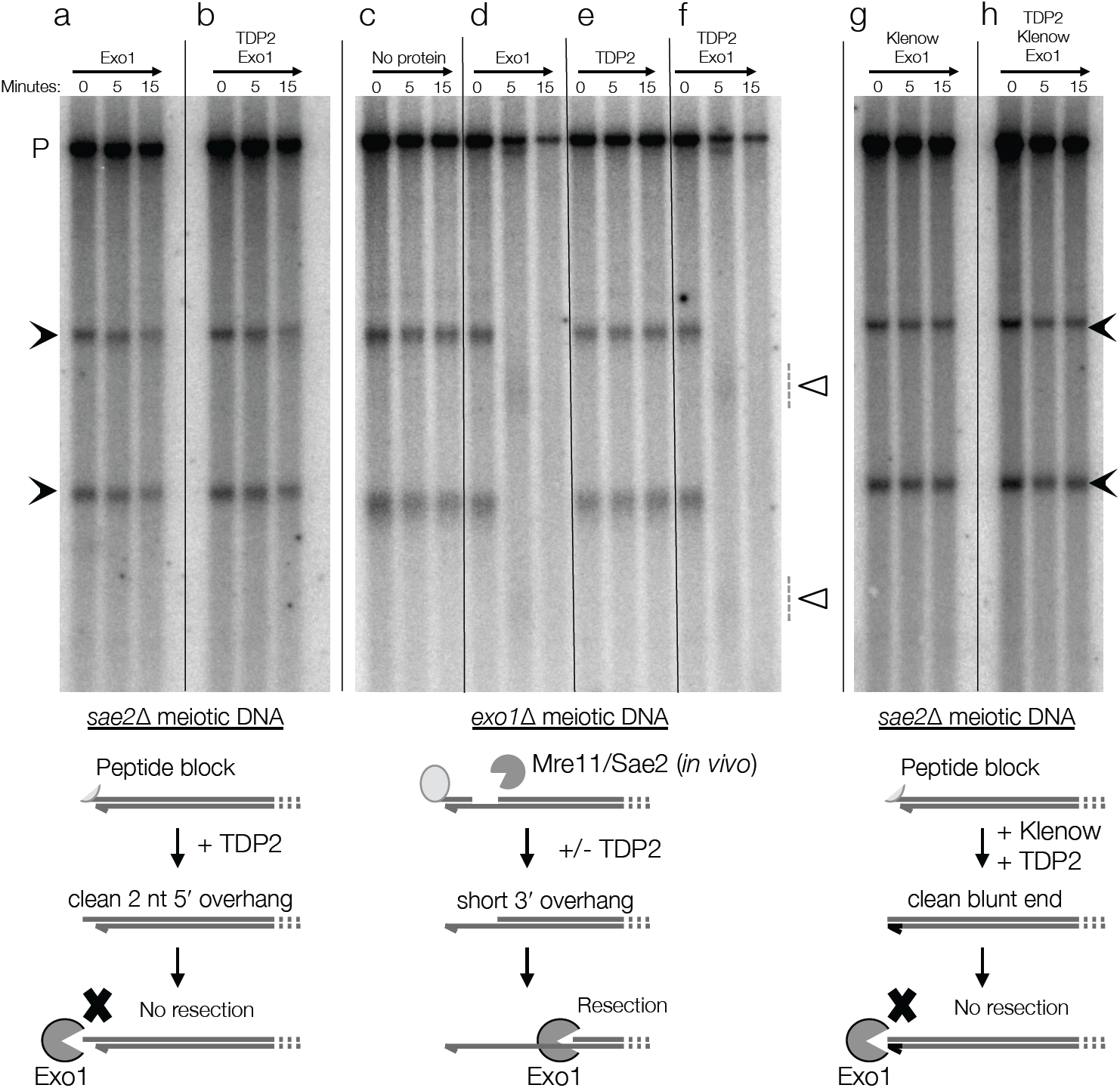
Recombinant Exo1 is unable to initiate 5′-3′ resection on 3′-recessed and blunt-ended Spo11-DSBs even when uncapped by recombinant TDP2, but can efficiently resect 5′-recessed Spo11-DSBs that have been processed by Mre11 *in vivo*. a-b, Genomic DNA isolated from meiotic *sae2*Δ cells was incubated with recombinant Exo1 without (a) or with (b) prior treatment with TDP2. c-f, Genomic DNA isolated from meiotic *exo1*Δ cells was incubated without (c), or with recombinant Exo1 (d), TDP2 (e), or TDP2 then Exo1 (f). c-d, As in (a, b), but TDP2-uncapped Spo11-DSB ends were blunted by treating with Klenow polymerase + dNTPs prior to incubation with Exo1. DNA was purified and digested with *Pst*I, separated by electrophoresis on a 0.7% agarose gel, blotted to nylon membrane and hybridised with a probe (*MXR2* locus) close to the *HIS4*∷*LEU2* meiotic recombination hotspot. The location of the two major Spo11-DSBs at this locus are marked with solid arrowheads. In *exo1*Δ cells, *in vivo* removal of Spo11 from DSB ends and partial resection causes the DSB signals to migrate slightly faster than in *sae2*Δ samples. Further smearing (open triangles) and subsequent disappearance of these signals following Exo1 treatment is due to rapid exonucleolytic degradation from the partially resected DSB ends. Lower panels are schematics of each set of reactions.

As a positive control to ensure that this inability to resect the Spo11-DSB end was not due to a problem with our recombinant Exo1 preparation, we repeated the same set of assays with genomic DNA isolated from an *exo1*Δ mutant (rather than *sae2*Δ). Under these conditions, MRX and Sae2-mediated nucleolytic cleavage of Spo11 from the DSB *in vivo* results in a short 3′ ssDNA tail (up to ∼300 bp) (15). As expected, incubation of this substrate with recombinant Exo1 resulted in specific degradation of the DSBs whether or not they were first treated with TDP2 (Fig 3c,d).

As a final control, we confirmed that TDP2 was active in the Exo1 buffer by incubating the Spo11-oligo substrate with TDP2 under the same conditions that the Exo1 resection assays were performed (Fig S1b). Collectively these results demonstrate that whilst TDP2 may have the necessary biochemical activity to remove covalently attached Spo11 from DSB ends, the resulting clean DNA ends are a poor substrate for Exo1 to initiate resection.

### Exo1 is unable to resect Spo11-DSBs that have been blunted by Klenow fragment

Spo11 creates a 2 nt 5′ overhang at the DSB ends (7, 39, 40). Prior *in vitro* assays using relatively short (∼3.0 kb) plasmid fragments indicated that the preferred substrate for Exo1 is a dsDNA end with a 3′ extension, similar in form to the resected DSBs created by MRX/Sae2, and in agreement with our observations presented above (41). By contrast, Exo1 displayed reduced, but not abolished, activity on substrates with a 4 nt 5′ extension, and only moderately reduced activity on a blunt ended substrate (41). These latter findings contrast somewhat with our observations that even just a 2 nt 5′ overhang completely prevents Exo1-catalysed resection of the TDP2-processed Spo11-DSBs (Fig 3a,b). A possible explanation for the difference may be the ratio of dsDNA to DSB ends, which is ∼100-fold greater in our assay system—and a much closer mimic of the conditions present *in vivo* during meiosis.

To determine whether Exo1 was able to resect blunt DSB ends under these same ‘*in vivo*-like’ conditions, we repeated the experiments utilising the TDP2-processed *sae2*Δ genomic DNA, but this time including a preincubation step with Klenow fragment and dNTPs to blunt the 2 nt 5′ overhang prior to incubation with Exo1 (Fig 3g,h). Unlike for the partially resected substrates, no resection by Exo1 was detectable even when DSB ends were first blunted by Klenow fragment (Fig 3g,h). Taken together our results suggest that Exo1 is acutely sensitive to the structure of the DNA end, with partially resected ends—such as those created by MRX/Sae2-dependent processing—being a significantly favoured substrate when excess competitor dsDNA substrate is present. We suggest that our observations may represent a closer match to the *in vivo* situation, where—even in meiosis—DSB ends are relatively infrequent compared to the intact double-stranded genomic DNA.

### Ectopic expression of TDP2 is unable to cleave Spo11 from DSB ends in *S. cerevisiae*

An orthologue of the TDP2 protein is not present in *S. cerevisiae*. However, human TDP2 was identified via its ability to rescue the sensitivity of yeast cells to camptothecin (a Top1 poison), and to also suppress etoposide sensitivity (a Top2 poison) indicating that human TDP2 is active when expressed in *S. cerevisiae* cells (25). Given our findings that TDP2 has the appropriate biochemical activity to remove Spo11 from both ssDNA and dsDNA *in vitro* (above), we were interested to investigate whether TDP2 expression during *S. cerevisiae* meiosis would permit removal of Spo11 *in vivo*.

To test this idea, we placed His-tagged TDP2 under the control of the *ADH1* promoter in a *sae2*Δ strain, and induced cells to enter meiosis (Fig 4a). Genomic DNA was harvested at hourly intervals during meiotic prophase, and used to probe the DSB hotspot at the *HIS4*∷*LEU2* locus as in earlier experiments (Fig 2 and Fig 3). We observed no difference in the abundance or electrophoretic mobility of the DSB bands in either the presence or absence of TDP2 expression, indicating that TDP2 was neither promoting resection nor repair of the Spo11-DSBs (Fig 4a). Expression of TDP2 from an inducible *GAL1* promoter construct after 4 hours in meiosis in a *sae2*Δ strain also resulted in no detectable alteration in DSB signal (data not shown).

**Figure 4:**
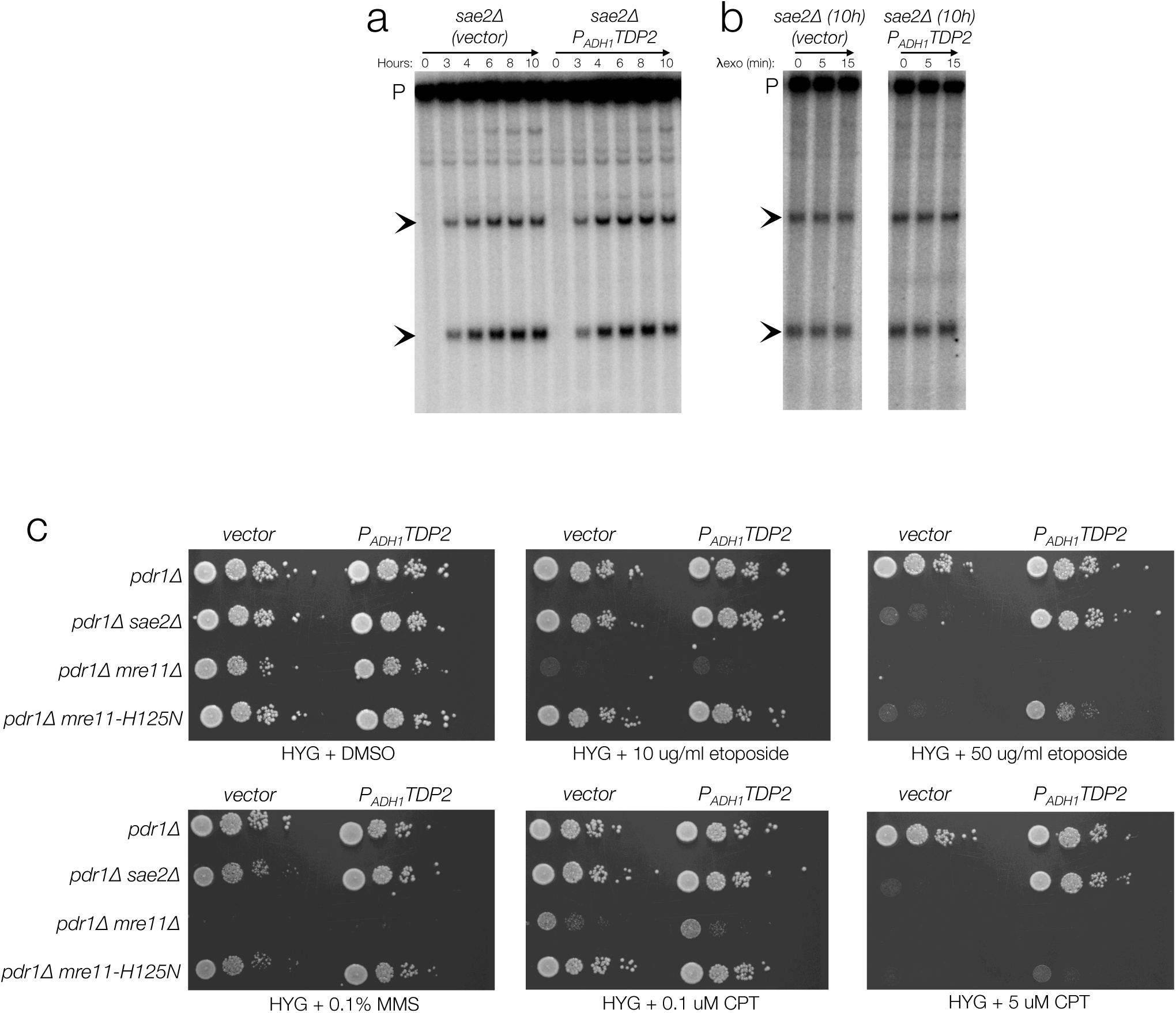
Ectopic expression of *TDP2* cannot remove Spo11 from DSB ends *in vivo*, but can suppress camptothecin and etoposide sensitivity. a-b, Genomic DNA was isolated at the indicated timepoints from a synchronous meiotic culture of *sae2*Δ cells harbouring either an empty vector, or one expressing *TDP2* from the *ADH1* promoter (*P*_*ADH1*_*TDP2*). DNA was purified and digested with *Pst*I, separated by electrophoresis on a 0.7% agarose gel, blotted to nylon membrane and hybridised with a probe (*MXR2* locus) close to the *HIS4*∷*LEU2* meiotic recombination hotspot. The location of the two major Spo11-DSBs at this locus are marked with solid arrowheads. In *sae2*Δ cells, Spo11 is not removed and resection cannot occur, causing DSB species to migrate as a tight band. Expression of TDP2 does not alter this migration pattern (a). b, DNA from the 10 h timepoints was incubated with lambda exonuclease for the indicated length of time. c, The indicated strains harbouring either an empty vector, or one expressing *TDP2* from the *ADH1* promoter (*P*_*ADH1*_*TDP2*) were grown to log phase, serial diluted (10-fold), and spotted onto plates containing 300 µg/ml hygromycin (HYG), to maintain plasmid selection, and the indicated concentrations of etoposide, camptothecin (CPT) or methyl methanesulphonate (MMS), and incubated for 3 days at 30°C.

A possible explanation for no visible resection or repair of Spo11-DSBs after TDP2 expression is that TDP2 has removed Spo11 from the DSB end, but these DSBs, with 2 nt 5′ overhangs, are resistant to nucleases such as Exo1 (as demonstrated *in vitro* in Fig 3), and that NHEJ components may be downregulated in meiosis perhaps due to increasing CDK activity as has been observed in cycling *S. pombe* cells (42). To test this possibility, we isolated genomic DNA from a 10 h meiotic time point from cells expressing *TDP2*, or from control cells, and incubated with lambda exonuclease (Fig 4b). At this time point, if TDP2 was capable of removing Spo11 from the 5′ end of the DSBs, lambda exonuclease would be able to resect these DSBs (as described in Fig 2). However, we detected no visible resection in the TDP2-expressing nor control cells (Fig 4b). These results suggest that despite the ability of TDP2 to suppress topoisomerase-induced DNA damage in *S. cerevisiae* (25), Spo11-DSBs are refractory to processing by the TDP2 enzyme.

### Ectopic expression of TDP2 rescues sensitivity of *sae2*Δ and *mre11-H125N* mutant cells to etoposide and camptothecin

As a final investigation of TDP2 activity *in vivo* in *S. cerevisiae* cells, we expressed TDP2 in drug-sensitive (*pdr1*Δ) haploid cells (43), and tested for the rescue of camptothecin (CPT) and etoposide sensitivity, as well as to the DNA-alkylating agent methyl methanesulphonate as a control (MMS). At the concentrations tested, wild type control cells (*pdr1*Δ) were not sensitive to etoposide or camptothecin, and there was no effect of TDP2 expression (Fig 4c). *sae2*Δ cells were sensitive to the higher concentration of etoposide and CPT, and were rescued efficiently by ectopic TDP2 expression (Fig 4c). By contrast, whilst loss of Mre11 nuclease activity (*mre11-H125N*) resulted in drug sensitivities that were similar to the *sae2*Δ mutant, TDP2 expression caused much weaker suppression than in the *sae2*Δ cells— particularly following CPT treatment (Fig 4c). These observations suggest a separable role for Sae2 and Mre11 in the repair of Top1- and Top2-induced DNA lesions, with TDP2 only able to fully complement the functions of the Sae2 protein. Cells bearing a full deletion of *MRE11* were highly sensitive to even the lower drug concentrations, with no rescue of growth observed upon expression of TDP2 (Fig 4c). This latter result is consistent with the inability for TDP2 to complement some or all of the many functions of Mre11 (44). Notably, TDP2 expression suppressed a very minor growth defect of *sae2*Δ and *mre11-H125N* cells in the presence of MMS (compare colony sizes in Fig 4c with and without TDP2 expression), suggesting either that MMS treatment leads to the accumulation of topoisomerase-lesions, or that it generates directly or indirectly, other potentially toxic intermediates that are substrates for the phosphodiesterase activity of TDP2.

## DISCUSSION

TDP2 was identified in a screen for mammalian genes that rescued the sensitivity of *S. cerevisiae* cells to the Top1 poison, camptothecin (25). Biochemical analyses subsequently demonstrated the preferred activity of TDP2 against synthetic 5′ phosphotyrosine groups (25), as well as larger peptides derived from partially proteolysed Top2 (31). Most recently, TDP2 was shown to be proficient in catalysing the release of intact Top2 from dsDNA ends in a reaction that is strongly dependent upon the ZATT protein (45). Here we have characterised the activity of TDP2 against the 5′-linked Spo11 complexes that arise during the induction of meiotic recombination. Recombinant human TDP2 displays robust removal of partially proteolysed Spo11 protein from both ssDNA oligos and dsDNA molecules derived from meiotic cell extracts. We find that non-proteolysed Spo11 is also released albeit with reduced efficiency.

Under normal conditions in meiosis, Spo11-DSBs are processed by the nuclease activities of Mre11 in conjunction with Sae2, followed by extensive 5′ to 3′ resection mediated by Exo1. Surprisingly, despite efficient *in vitro* removal of Spo11 from DSB ends by TDP2, we have found that the resulting protein-free DNA ends remain very poor substrates for Exo1, in contrast to DSB substrates containing a terminal 3′ single-strand region of ∼300 nt. These observations agree with prior observations indicating a preference of Exo1 for partially single-stranded substrates (12, 41). However, our experiments are the first that employ physiological frequencies of DSB ends relative to genome content (approx. 1 DSB end for every 150 kb), and lead us to conclude that excess dsDNA acts as a competitive inhibitor of Exo1’s ability to initiate resection at clean, relatively blunt, dsDNA ends—but not at dsDNA ends that have already undergone limited resection.

Such competitive inhibition may also influence the activity of other proteins involved in DSB end resection. For example, the Sae2-stimulated cleavage of protein-linked dsDNA by Mre11, has thus far only been demonstrated on short, synthetic dsDNA substrates (11–13). Attempts to reconstitute a similar Mre11 endonuclease reaction on natural Spo11-DSBs isolated from meiotic cells has thus far proved unsuccessful in our laboratory (D. Johnson and M. Neale, unpub. obs.)—perhaps in agreement with the reduced efficiency of the Mre11 endonuclease reaction on longer synthetic substrates (Ref 13 and E. Canavo and P. Cejka, unpub. obs.). It is interesting to note that in *S. cerevisiae* Mre11 is loaded at meiotic recombination hotspots prior to Spo11-DSB formation (46). It is tempting to speculate that this interaction is important to ensure efficient nucleolytic processing of Spo11-DSB ends in *S. cerevisiae*—thereby ensuring that meiotic DSB repair is directed down the HR repair pathway.

In *C. elegans*, loss of COM-1 function (the Sae2/CtIP orthologue) leads to aberrant DSB repair and meiotic chromosome segregation due to activation of the NHEJ repair pathway (21, 22). Such observations suggest that mechanisms other than Mre11 are competent to release Spo11 from meiotic DSB ends, generating ligatable DNA ends—at least under conditions when the preferred Mre11-pathway is disrupted. An orthologue of human TDP2 exists in worms (33), but it is yet to be determined whether this activity is the one responsible for enabling NHEJ repair of Spo11-DSBs in the *com-1* mutant.

Interestingly, whilst human TDP2 is capable of catalysing Spo11 removal from DSB ends *in vitro*, we were unable to detect Spo11 removal *in vivo* following ectopic expression of TDP2 within meiotic *S. cerevisiae* cells. Whilst it is hard to rule out a technical reason for observing this negative result, we *were* able to observe robust rescue of camptothecin and etoposide sensitivity in *sae2*Δ and *mre11*-‘nuclease-dead’ mutant strains upon TDP2 expression. Thus, our favoured interpretation is that Spo11-DSB ends in *S. cerevisiae*—unlike topoisomerase-DNA lesions—are inaccessible to the TDP2 phosphodiesterase activity. Such inaccessibility may be due to a higher order protein complex present at meiotic DSBs, which are known to require directly or indirectly at least a dozen independent proteins (47). Such ideas are also consistent with observations suggesting multimerisation of the Spo11 enzyme within recombination hotspots (D. Johnson and M. Neale, manuscript in preparation). Alternatively, or in addition, topoisomerase-induced lesions may be subject to greater rates of proteolysis and/or denaturation than Spo11, thereby stimulating the end-cleaning activity of TDP2.

Recognition, and appropriate repair, of DNA lesions is critical for cell survival. During meiotic prophase, the channelling of Spo11-DSBs down the HR pathway has further importance to enable crossover formation, reductional chromosome segregation, and for the generation of genetic diversity (48). Our observations here provide insight into the mechanisms of repairing such protein-linked DSB ends created by topoisomerase-like enzymes such as Spo11. Looking more broadly, our findings may also help to explain some of the genetic differences observed for the resection and repair of ‘clean’-ended site-specific DSBs created by nucleases versus the complex DSB ends created by ionising radiation, which may be a mixture not just of DNA ends with base damage, but also those with blunt or partial 5′ extensions, or occluded by larger protein moieties.

## ACKNOWLEDGEMENTS

We thank E. Hoffmann, N. Hunter, S. Keeney and J. Nitiss for yeast strains; L. Symington for plasmids; K Caldecott for recombinant human TDP2 protein. D.J. is supported by an ERC Consolidator Grant (‘DNA-Repair-Chromatin’) to M.J.N. M.J.N. was supported by this award and a University Research Fellowship from the Royal Society, and a Career Development Award from the Human Frontiers Science Program Organisation.

## AUTHOR CONTRIBUTIONS

D.J. and M.J.N. designed and performed the experiments and wrote the paper. E.C. and P.C provided essential reagents and technical supervision.

## METHODS

### Strains, Plasmids and Recombinant Proteins

Meiotic experiments utilised the *S. cerevisiae* SK1 background (49), and harboured the *sae2*Δ∷kanMX or *exo1*Δ∷kanMX full gene deletions (9). For Spo11-oligo enrichment, strains harboured the Spo11-FLAG_3_His_6_∷kanMX construct (30). The multicopy constitutive *P*_*ADH1*_*TDP2* plasmid (pDJ85) was prepared after replacing the *URA3* gene in pVTU260TT (25) with a Hygromycin resistance cassette. Cell viability spot tests utilized the S288c background harbouring the dominant multi-drug sensitizing cassette *pdr1*Δ∷*pdr1DBD*-*CYC8* (referred to as ‘*pdr1*Δ’) (43) and additionally harbouring the *sae2*Δ∷kanMX, *mre11*Δ∷*URA3* gene deletions or *mre11-H125N* point mutation (50). Genetic modifications were introduced into yeast strains using standard procedures (51). Recombinant human His-TDP2 was a gift from K. Caldecott, and prepared as described previously (25). Recombinant *S. cerevisiae* Exo1-FLAG and RPA was prepared as described previously (41).

### Isolation and analysis of meiotic DNA breaks

SK1 cells were induced synchronously into meiosis using standard procedures (52). At indicated timepoints, 10 ml of cell culture was spheroplasted in 1M sorbitol / 50 mM EDTA / 50 mM NaHPO_4_ buffer, and lysed by addition of SDS to 0.5% in the presence of proteinase K at 60°C. Nucleic acids were purified by phenol extraction and ethanol precipitation, and resuspended in 1×TE. DSB signals at the *HIS4*∷*LEU2* recombination hotspot were detected using standard techniques (53). Briefly DNA was digested with *Pst*I, electrophoresed on 0.7% agarose in 1×TAE for approximately 18 h at room temperature, transferred to nylon membrane under denaturing conditions then hybridized with a probe that recognizes the *MXR2* locus. Prior to digestion, as appropriate, samples were equilibrated in reaction buffer (25 mM HEPES, pH 8.0, 130 mM KCl, 1 mM DTT, 10 mM MgCl2) and incubated with 300 nM recombinant TDP2 for 1 hour at 37°C, 5 units of lambda exonuclease (NEB) for 1 hour at 37°C, 15 units of Klenow (3′→5 ′ exo–) polymerase (NEB) with 66 nM dNTPs for 30 minutes at 30°C, 20 mM recombinant Exo1-FLAG with 680 nM RPA (REF) for 30 minutes at 30°C, or combinations thereof as indicated in each figure. Reactions involving Exo1 were performed in a similar buffer (25 mM Tris acetate, pH 7.5, 1 mM DTT, 5 mM magnesium acetate, 100 mM sodium acetate, 0.25 mg/mL BSA) in which TDP2 maintains activity (Figure S1b). Reactions were stopped and DNA extracted by the addition of 1 volume of water and 1 volume of phenol:chloroform:isoamyl alcohol (25:24:1) prior to restriction digestion and electrophoresis. Radioactive signals were collected on phosphor screens, scanned with a Fuji FLA5100 and quantified using ImageGauge software.

### Isolation and analysis of Spo11-oligo complexes

Spo11-oligonucleotide complexes were detected by immunoprecipitation and end-labelling following established methods (54). Briefly, cells were broken in 10% ice-cold TCA using zirconium beads and a BioSpec 24. Precipitated material was dissolved in SDS buffer, diluted with Triton X100, and Spo11 was immunoprecipitated from total soluble protein using anti-FLAG antibody (Sigma) and protein-G-agarose (Roche). Oligonucleotide complexes were labelled with alpha-^32^P dCTP using terminal deoxynucleotidyl transferase (Fermentas) and fractionated on a 7.5% SDS-PAGE gel. Alternatively, radiolabelled Spo11-oligo complexes were precipitated with 90% ethanol at –80°C overnight without or following digestion with trypsin or proteinase K (Fisher) for 1 hour at 37°C. Precipitated material was collected by centrifugation, solublised in reaction buffer (25 mM HEPES, pH 8.0, 130 mM KCl, 1 mM DTT, 10 mM MgCl2), and incubated with the stated concentration of recombinant TDP2 for 30 minutes at 30°C prior to denaturation in formamide loading dye and separation on 19% urea/PAGE/1×TBE. Radioactive signals were collected on phosphor screens, scanned with a Fuji FLA5100 and quantified using ImageGauge software.

### Cell viability spot tests

YPD cultures were grown in the presence of 300 µg ml^-1^ hygromycin (Invitrogen) to maintain selection of the plasmids. Overnight cultures were diluted 40-fold in fresh YPD containing 300 µg ml^-1^ hygromycin and grown for 4 hours. Cultures were diluted to 0.2 OD_600_ and a 10-fold dilution series prepared before spotting out 3 µl onto the YPD plates containing 300 µg ml^-1^ hygromycin and the stated drug dose or DMSO solvent control. Plates were incubated at 30 °C for 3 days prior to image acquisition.

**Table 1.**
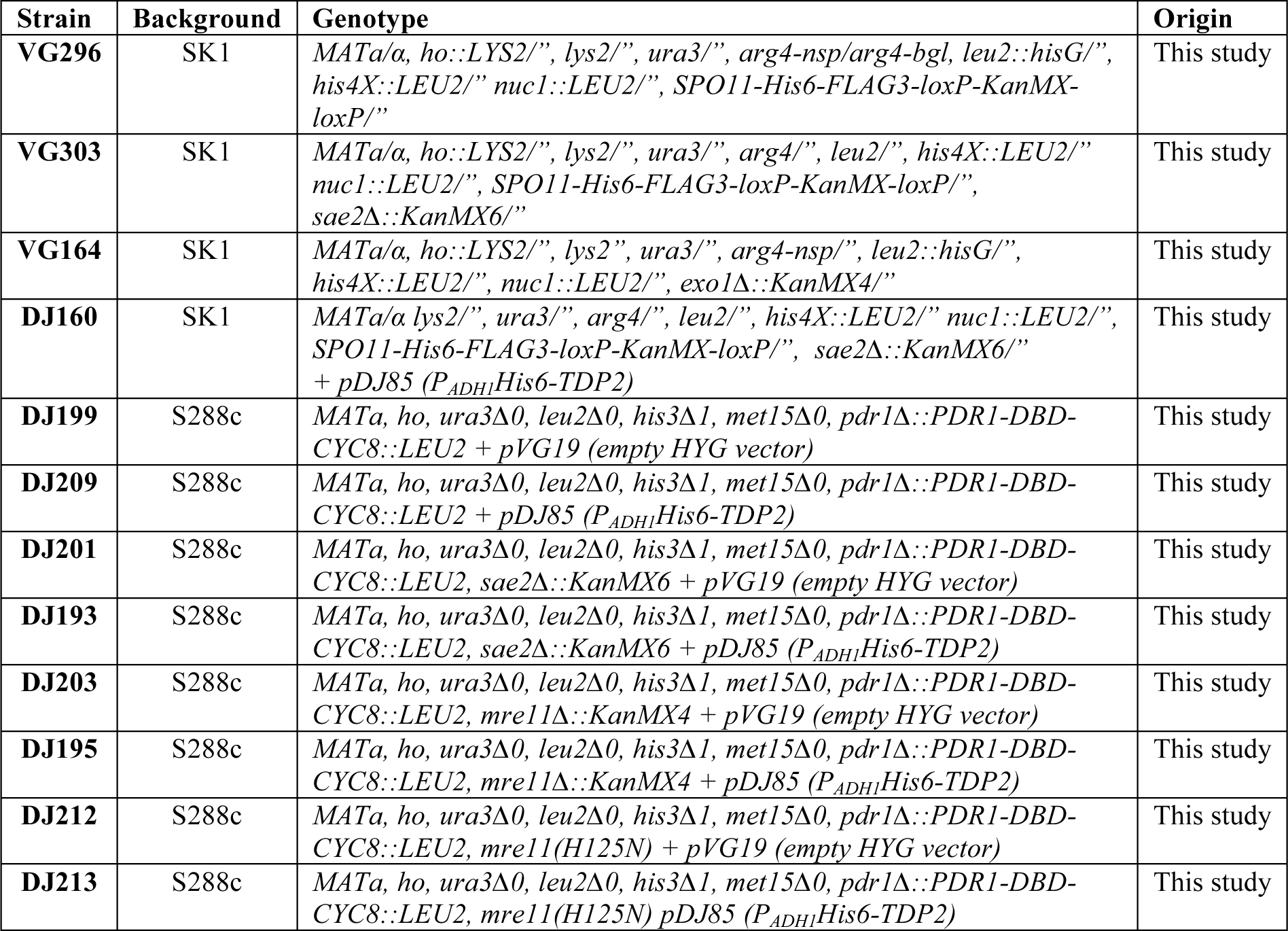
*S. cerevisiae* strains used in this study.

**Figure 5:**
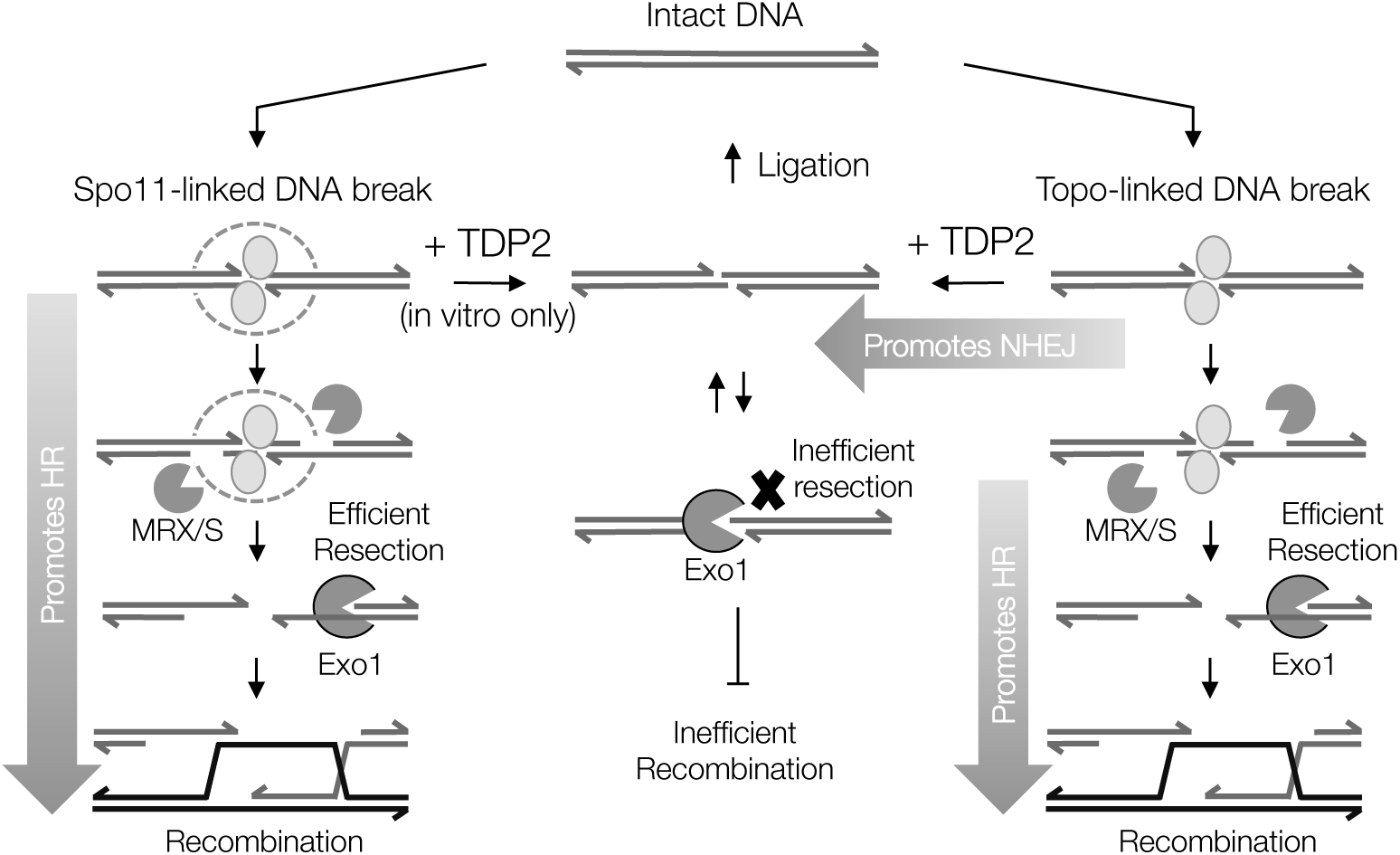
Model of DSB repair via HR or NHEJ depending on repair mechanism. Covalently bound protein at DNA ends has to be removed in order for repair to occur. Mre11 and Sae2 remove the protein block (Spo11 or topoisomerases) nucleolytically, generating a 3′ ssDNA overhang substrate that is refractory to NHEJ, but the preferred substrate for long-range resection by Exo1, thereby promoting homologous recombination (left and right panels). Removal of covalently bound protein, for example, by hydrolytically cleaving the phosphotyrosyl bond between the protein and the 5′ end of the DSB (e.g. by TDP2) generates clean, short 5′ overhang ends that are refractory to resection by Exo1—thus potentially inhibiting repair by HR—but the complementary nature of the two ends may promote repair by NHEJ, for example at Top2-DSBs (central panel). The inability for ectopic expression of TDP2 to suppress the requirement for Sae2 in S. *cerevisiae* meiosis suggests that Spo11-DSBs, unlike Topoisomerase-DNA breaks, are inaccessible to TDP2, perhaps due to a large complex at the Spo11-DSB site occluding access to TDP2 (dotted circle in left panel).

---

**Figure S1:**
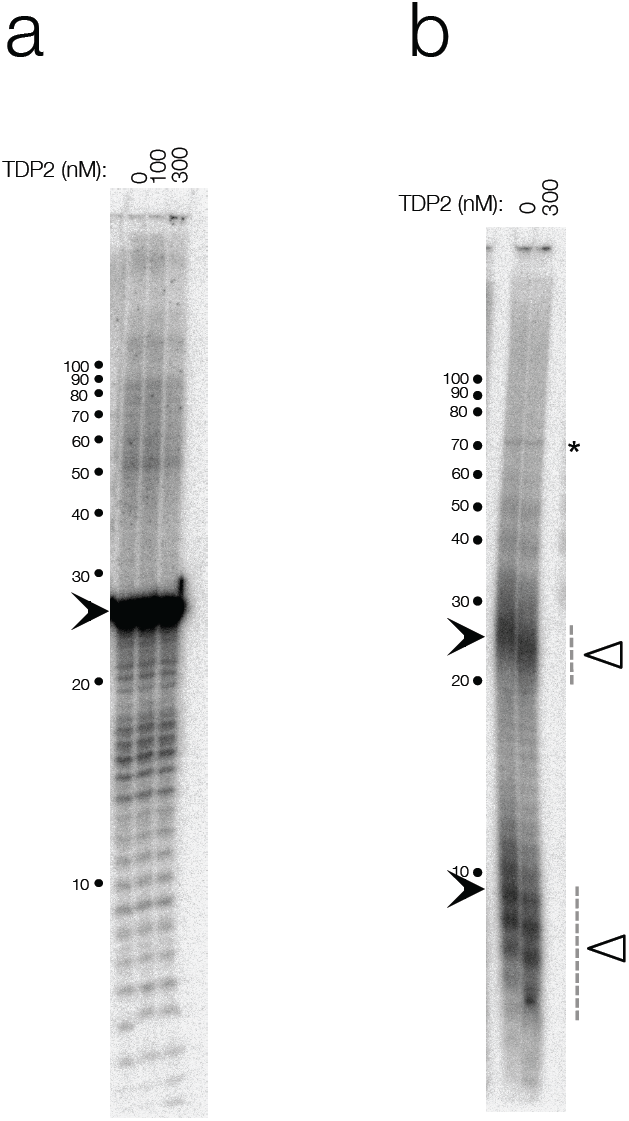
Recombinant TDP2 is free of nuclease activity, and is active in the preferred Exo1 reaction buffer. a, To test for trace nuclease activity within the recombinant TDP2 preparation, a 30 nt oligonucleotide of random sequence was 3′-labelled using TdT and radiolabelled dCTP then incubated with the stated concentrations of TDP2 at 37 °C for 1 hour then separated by denaturing urea-PAGE (19% acrylamide). The 30 nt oligo is indicated with a filled arrowhead. **b,** Proteinase K-treated 3′-labelled Spo11-oligo complexes as in Fig 1f were incubated with the indicated concentration of TDP2 at 37°C for 1 hour in Exo1 reaction buffer (25 mM Tris acetate, pH 7.5, 1 mM DTT, 5 mM magnesium acetate, 100 mM sodium acetate, 0.25 mg/mL BSA) then separated by denaturing urea-PAGE (19% acrylamide). Solid arrowheads indicate the migration position of Spo11-oligo species (prior to TDP2 treatment). Open triangles indicate the migration position of Spo11-oligos after Spo11-peptide removal by TDP2.

